# A strategy to optimize the peptide-based inhibitors against different mutants of the spike protein of SARS-CoV-2

**DOI:** 10.1101/2022.02.27.482153

**Authors:** Prerna Priya, Abdul Basit, Pradipta Bandyopadhyay

## Abstract

SARS-CoV-2 virus has caused high-priority health concerns at a global level. Vaccines have stalled the proliferation of viruses to some extent. Yet, the emergence of newer, potentially more infectious, and dangerous mutants such as delta and omicron are among the major challenges in finding a more permanent solution for this pandemic. The effectiveness of antivirals Molnupiravir and Paxlovid, authorized for emergency use by the FDA, are yet to be assessed at larger populations. Patients with a high risk of disease progression or hospitalization have received treatment with a combination of antibodies (antibody-cocktail). Most of the mutations leading to the new lineage of SARS-CoV-2 are found in the spike protein of this virus that plays a key role in facilitating host entry. The current study has investigated how to modify a promising peptide-based inhibitor of spike protein, LCB3, against common mutations in the target protein so that it retains its efficacy against the spike protein. LCB3 being a prototype for protein-based inhibitors is an ideal testing system to learn about protein-based inhibitors. Two common mutations N501Y and K417N are considered in this work. Using a structure-based approach that considers free energy decomposition of residues, distance, and the interactions between amino acids, we propose the substitutions of amino acid residues of LCB3 inhibitors. Our binding free energy calculations suggest a possible improvement in the binding affinity of existing inhibitor LCB3 to the mutant forms of the S-protein using simple substitutions at specific positions of the inhibitor. This approach, being general, can be used in different inhibitors and other mutations and help in fighting against SARS-CoV-2.

## 1. Introduction

SARS-CoV-2 is a novel, positive-sense, single-stranded RNA virus, belonging to the family Coronaviridae that emerged at the tail end of 2019 ^1^. It primarily affects the respiratory system ranging from mild to severe infection ^2^. This virus is highly infectious and can easily transmit from one person to another ^3^. This is responsible for the sudden outbreak of a pandemic that collapsed the global health care system and economy. At least 43 crores of people have been infected with SARS-CoV-2 and among them, more than 59 lakhs died (https://covid19.who.int/, data as of Feb 25, 2022). This crisis prompted scientists, doctors, manufacturers, and regulating authorities to race against time and develop treatments as well as vaccines.

SARS-CoV-2 consists of ~30 kb nucleotides which encode approximately 29 proteins including four structural proteins (spike, membrane, envelope and nucleoprotein) ^4–5^. The potential therapeutic targets of this virus include spike (S), envelope (E), membrane (M), nucleoprotein (N), replicase polyprotein (chymotrypsin-like protease (3CLpro), or main protease (Mpro), papain-like protease (PLpro), RNA dependent RNA polymerase (RdRp), and other non-structural proteins) and transmembrane serine protease 2 (TMPRSS2) ^6–7^ Strategies used to develop inhibitors against these potential therapeutic targets include i) drug repurposing ^8^ either by experiment-based high throughput screening or structure-based virtual screening, ii) fragment-based drug designing ^9^, or iii) *de novo* drug design ^10^. Spike protein and TMPRSS2 play a crucial role in the entry of viruses therefore the therapeutics designed against this protein can prevent the viral entry. Several peptide-based inhibitors were designed and tested against the S-protein of SARS-CoV-2 ^11–12^. Hydrophilic compound, ‘Salvianolic acid C’ obtained from traditional Chinese medicine potently inhibits the membrane fusion of S-protein^13^. Antiviral ‘arbidol’ inhibits the trimerization of S-protein, however, no significant benefit of this molecule was observed in the clinical trial ^14–17^. Novel drug-like compounds DRI-C23041 and DRI-C91005 inhibited the interaction of hACE-2 with S-protein in cell-free ELISA assay ^18^. Chen et al. discovered six inhibitors against S-protein (Cepharanthine, abemacicilib, osimertinib, trimipramine, colforsin, and ingenol) that were tested against pseudotyped particles SARS-S and MERS-S ^19^. MM3122, camostat mesylate, nafamostat, MNP10 (marine natural product 10) are some of the potential inhibitors of TMPRSS2 ^20–27^. The E-protein of SARS-CoV-2 consists of 75 residues that form a homopentameric cation channel and are involved in the viral assembly, budding formation of the envelope and pathogenesis ^28–29^. Amantadine, Rimantadine, Hexamethylene amiloride (HMA) and several flavonoids act as potent inhibitors against E-protein ^28, 30^. Several known inhibitors (e.g. nelfinavir, lopinavir, ritonavir) have been tested and show high binding affinity against the 3CLpro or Mpro that play a crucial role in the proteolytic processing of replicase polyprotein ^31–34^. Being involved in the replication and transcription of the SARS-CoV-2 genome, RdRp is a potential drug target for SARS-CoV-2 ^35^. Nucleotide analog Remdisivir and favipiravir bind efficiently with RdRp ^36–38^. PLpro is a proteolytic enzyme and also affects the host antiviral immune response that leads to the viral spread ^39–40^. GRL0617 is a promising inhibitor against PLpro^41^. Emergency use authorization has been given to several vaccines, monoclonal antibodies as well as antiviral drugs developed by various pharmaceutical companies to save lives during the pandemic ^42–45^. As a result, more than 10 billion doses of vaccine have been already administered to curb the progress of COVID-19 (https://covid19.who.int/). Remdesivir, an antiviral drug, was used widely but did not show significant clinical benefit against COVID-19 ^46^. Monoclonal antibodies ‘sotrovimab’ as well as the antibody cocktail - a combination of ‘casirivimab’ and ‘imdevimab’ have shown clinical benefits and reduced the hospitalization rate in patients with COVID-19 (https://www.covid19treatmentguidelines.nih.gov/) ^45, 47-48^. However, none of these has offered a real cure against this disease yet. People are also getting infected, post-vaccination, with new variants though showing milder infection ^49^. Recently, FDA authorized two COVID-19 antiviral pills: ‘Molnupiravir’ (Merck, USA) and, ‘Paxlovid’ (Pfizer, USA) for emergency use in patients who are at high risk ^50–52^. Though these pills worked well in clinical trials, the real-world efficacies of these pills are yet to be assessed.

One of the major challenges of treating COVID-19 is the appearance of different mutant strains of the virus. The mutation frequency of SARS-CoV-2 to form newer strains such as alpha, beta, a more aggressive delta, and the most recent omicron, is a major challenge in developing a treatment for COVID^53-56^. It also includes the question, of whether the treatments developed for SARS-CoV-2 will work on its existing and upcoming variants^57–59^. The majority of the mutations reported are in the spike (S) glycoprotein, which is responsible for the entry of the virus inside the human host via human receptor protein Angiotensin Converting Enzyme-2 (ACE-2)^60–62^. In this work, we want to check if existing inhibitors can be modified in such a way that these retain their efficacy for the mutant forms. For a proof-of-principle study, we have taken a designed mini-protein inhibitor, LCB3, a prototype for protein-based inhibitors including antibodies ^12^ (Figure 1). LCB3 is a 64 amino-acid-long, stable, potent miniprotein inhibitor that competes with ACE-2 and binds tighter with the S-protein ^12^. It has the potential to be used as a therapeutic and may open options for direct delivery to the nasal passage or other parts of the respiratory system. However, several studies have shown mutations induced alteration in the binding affinity of S-protein to ACE-2 receptor^63-66^. S_N501Y_ (Spike protein of SARS-CoV-2 with N501Y mutation) binds to ACE-2 receptor with 7-fold higher affinity than the WT whereas S_K417N_ (Spike protein of SARS-CoV-2 with K417N mutation) binds with 4-fold lower affinity ^67^. This reflects the possibility of alteration in the binding affinity of the LCB3 inhibitors against the mutated S-protein and needs further investigation. Two common mutations, N501Y and K417N are present at the binding interface of the protein (Figure 1) were tested. We have investigated the effect of these two mutations on the binding affinity between LCB3 and S-protein and how to improve the binding affinity by modifying the inhibitors. N501Y mutation is found in various lineages including B.1.1.7 (alpha), B.1.351 (beta), and P.1 (gamma) variants first detected in the U.K., South Africa and Brazil respectively ^68^. K417N mutation is present in (beta) lineage B.1.351 and B.1.617.2 (delta plus-a sub-lineage of delta variant) ^69^. Both of these mutations (N510Y and K417N) were also reported in the newly detected variant of concern, lineage B.1.1.529 (omicron) ^70^. These mutations have immune evasion properties^71–73^.

**Figure 1:**
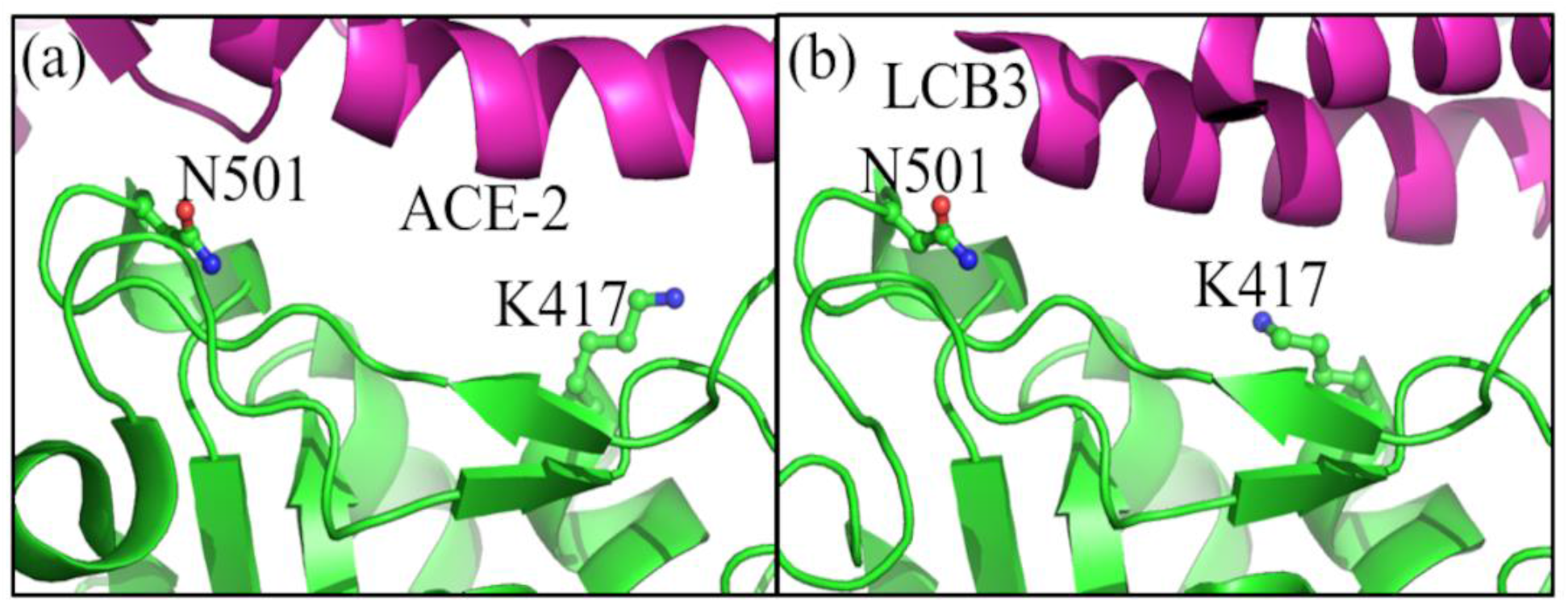
Binding Interface of spike protein (green) with (a) ACE-2 receptor (magenta) using PDB ID: 6M0J and (b) LCB3 miniprotein inhibitor (magenta) using PDB ID: 7JZM. K417 and N501 are shown in the stick.

To investigate the potential change in the binding affinity, associated with the mutations N501Y and K417N of S-protein with LCB3 and to improve the LCB3 inhibitor to make it effective against WT as well as mutants, we have used binding free energy calculation with the well-known Molecular Mechanics Poisson-Boltzmann Surface Area (MM-PBSA) method. We have decomposed the results of the binding affinity per residue of the inhibitor to see which residues contribute most to the binding. Then residues of LCB3 that are responsible for less binding affinity are changed to other residues based on the type of interaction and binding site geometry. This simple procedure increases the binding affinity of modified LCB3 with the two mutant forms of the proteins. This procedure is general and can be used to optimize other inhibitors against other mutations as well.

## 2. Materials and Methods

### 2.1 Structure Preparation

The coordinates of S-protein of SARS-CoV-2 with LCB3 inhibitor were retrieved from the protein data bank (PDB) with ID: 7JZM ^12^. This structure is used to model the N501Y and K417N mutant of the S-protein and variants of LCB3.

### 2.2 MD Simulation

ff14SB force field of AMBER 16 package was used to generate the parameters of the proteins^74–75^. All systems were solvated using TIP3P water molecules in a rectangular box using a minimum 10 Å distance between the edges of the box to the surface of the protein^76–77^. Special care was taken to preserve the disulfide bonds present between the cysteine residues present in PDB. Four pairs of Cysteines (C_336_-C_361_, C_379_-C_432_, C_480_-C_488_, C_391_-C_525_) of the S protein were involved in the disulfide bonding. Counter ions were added to neutralize the system and to maintain the 0.15M KCl salt concentration. 10000 steps of steepest descent followed by 10000 conjugant gradient minimization was used to remove the bad contacts in the solvated system ^78–79^. The systems were heated slowly up to 298 K for 50 ps followed by 50 ps density equilibration with the restraint weight of 2 kcal/mol/Å^2^ followed by and 500 ps equilibration run. After equilibration 10 independent simulations of all the complex, each of 10 ns was performed at NPT (by maintaining 300K temperature and 1 atmospheric pressure) to increase the sampling space (Table S1). For entropy calculations, separate long simulations (1 μs) of complex, receptor and ligands were performed (Table S2). Hydrogen bonds were constrained using a shake algorithm with 2 fs time integration^80^. The temperature was regulated by Langevin Dynamics with 2 ps of relaxation time^81^. 1 bar pressure was controlled by Berendesen’s barostat and periodic boundary conditions were applied for all the systems^82^. The particle mesh Ewald summation method was used for the calculation of the long-range electrostatic^83^. CPPTRAJ was used for further analysis of the trajectories obtained from simulation^84^.

### 2.3 Binding free Energy

MM-PBSA method was used to calculate the binding free energy of WT and mutants of S-protein with LCB3 and its variants. The single trajectory protocol for MM-PBSA was performed where the simulation with only the complex was performed, and from that simulation, properties of complex, receptor, and ligand are extracted. In this method, the standard free energy of binding (ΔG) can be defined by equation 1 which is implemented the in MMPBSA.py script of AMBER16 ^85^.

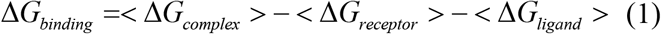

Here, ΔG_complex_, ΔG_receptor_, ΔG_ligand_ are the free energy of solvation for complex receptor, and ligand respectively. < > denotes the ensemble average.

ΔG_binding_ is calculated by equation 2:

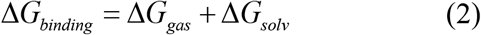

ΔG_gas_ is the change in the gaseous state. It is calculated by equation 5.

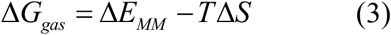

ΔE_MM_ includes electrostatic (ele) and van der Waals (vdW) components. Another component - TΔS represents the entropic contribution where T represents the temperature in Kelvin and S is the solute entropy.

ΔG_solv_ of equation 2 constitutes the polar and nonpolar components, (equation 4).

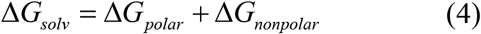

ΔG_polar_ (ΔG_PB_) is obtained from the solution of the Poisson Boltzmann equation. ΔG_nonpolar_ (ΔG_np_) constitutes of cavitation and dispersion terms (equation 5), where cavitation energy is estimated using a linear relation to the surface of the molecule and the dispersion is calculated using solute-solvent interactions^86^.

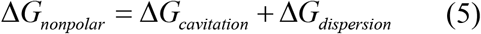

The nonpolar component is calculated using equation 6.

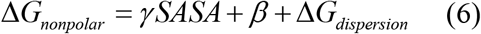

The value of γ and β is 0.0378 kcal/mol-Å^2^ and −0.569 kcal/mol respectively as implemented in the AMBER package. The solvent-accessible surface area (SASA) is used to distinguish the exposed and buried area of the protein. For that, a probe sphere with vdW radii of ~1.4 Å is rolled along the surface and if the probe can cross the area it is considered as surface accessible else the region is considered as buried where the solvent cannot enter.

The 0.15M ionic concentration was used for MM-PBSA calculations. The dielectric constant of the solute is highly dependent on the characteristics of the investigated system ^87–88^. In this study, various dielectric constants of solute were tested and the selection was done based on the similarity with the available data (details given in supporting information, Table S3). The dielectric constant of solute was kept 8. The total binding energy was averaged using 1000 frames obtained from 10 independent trajectories, each of 10 ns, for all the systems.

The entropy of the complex, receptor and ligand were estimated using Quasi Harmonic (QH) approximation method using CPPTRAJ. A sufficient phase space sampling is required for the reliable estimation of entropic contributions ^89^. For that, an overall 11 μs simulation was performed (Table S2). To estimate the effects of mutation on the flexibility of the system, entropy change (TΔS) was calculated using 5 lakhs frames from the separate 1 μs simulation, of complex, receptor and ligand.

## 3. Results and Discussions

### 3.1. Binding Energy Difference between WT and Mutant of S-protein with LCB3 inhibitor

The binding free energies of S_WT_ (wild-type S-protein), S_N501Y_ (asparagine at 501 position of S-protein mutated with tyrosine) and S_K417N_ (lysine at 417 position of S-protein mutated with asparagine) with LCB3 were determined using the MM-PBSA calculations and the values were compared. The binding affinity of S_K417N_ and S_N501Y_ mutants were reduced as compared with S_WT_ by ~ +8 kcal/mol (Table 1).

**Table 1:** The Binding free energy components of S_WT_, S_N501Y_ and S_K417N_, mutants of S-protein of SARS-CoV-2 with LCB3. The energy values are in kcal/mol. The calculations were averaged over 1000 frames obtained from 10 independent trajectories, each of 10 ns. The standard error is given in the parenthesis.

| System | $\Delta E_{ele}$ | $\Delta E_{vdW}$ | $\Delta G_{np}$ | $\Delta G_{PB}$ | $\Delta G_{ele + PB}$ | $\Delta G_{Total}$ |
| --- | --- | --- | --- | --- | --- | --- |
| $S_{WT}$ | -62.21(0.4) | 44.98 (0.7) | -66.42(0.4) | 59.80(0.3) | -2.41(0.4) | -23.85(0.6) |
| $S_{N501Y}$ | -49.77(0.6) | 43.75(1.0) | -59.98(0.6) | 50.31(0.5) | 0.54(0.6) | -15.69(0.6) |
| $S_{K417N}$ | -33.01(0.5) | 42.18 (0.9) | -61.41 (0.5) | 36.61(0.5) | 3.60(0.5) | -15.63(0.6) |
| $S_{N501Y} - S_{WT}$ | +12.44 | -1.23 | +6.44 | -9.49 | +2.95 | +8.16 |
| $S_{K417N} - S_{WT}$ | +29.2 | -2.80 | +5.01 | -23.19 | +6.01 | +8.22 |

The loss of +7.1 kcal/mol binding affinity of N501Y mutant of spike protein from WT to the mutant with LCB3 is reported by Williams et al. reflecting the correctness of our calculation protocol. ^90^ The S_N501Y_ mutation results in a loss of binding due to nonpolar solvation interaction (+6.44 kcal/mol). There is also a loss of +2.95 kcal/mol in polar contributions accompanied by the gain in the van der Waals (vdW) component by −1.23 kcal/mol (Table 1).

The lower binding affinity of S_K417N_ mutant is mainly due to the loss in polar interactions (+6.01 kcal/mol) and non-polar solvation (~+5.01 kcal/mol) with the slight gain of −2.80 kcal/mol of vdW interactions. It reflects that when the positively charged residue lysine was mutated with the uncharged polar asparagine residue, the polar interactions reduced.

### 3.2 Residue-Wise Free Energy Decomposition

The reduced binding affinity of LCB3 with S_K417N_ and S_N501Y_ mutants provided the scope to improve the LCB3 inhibitor further in such a way so that it can work potently and inhibit the WT as well as mutants. For that, residue-wise free energy decomposition was performed for the identification of important residues ^91^. Free energy contributions of each residue were decomposed into electrostatic, vdW, and polar solvation and the amino residues of LCB3 near the mutated residue of S-protein were analyzed (Table 2). D3 (aspartic acid at 3rd position) and E4 (glutamic acid at 4th position) of LCB3 are present near N501 (asparagine at position 501) of S-protein whereas T10 (threonine at 10th position) and D11 (aspartic acid at 11th position) of LCB3 present in the vicinity of K417 (lysine at 417 position) (Figure 2). There is the loss of+1.0 kcal/mol and +0.5 kcal/mol in the vdW component of D3 of LCB3 with S_N501Y_ and S_K417N_ respectively. These losses were compensated with the gain in the polar contributions, and the contribution of D3 was almost similar in S_N501Y_ whereas slightly lesser (+0.3 kcal/mol) in S_K417N_ (Table 2). However, the mutation (S_N501Y_ or S_K417N_) leads to the overall loss of ~+8 kcal/mol (Table 1). The loss in the energy contributions of E4 (+1.3 kcal/mol in S_N501Y_ and +0.5 kcal/mol in S_K417N_), T10 (+0.2 kcal/mol in S_N501Y_ and +0.7 kcal/mol in S_K417N_) and D11 (+0.9 kcal/mol in S_N501Y_ and +0.5 kcal/mol in S_K417N_) were observed in both S_N501Y_ and S_K417N_ (Table 2). It is likely that the interactions between these residues of LCB3 and neighboring residues of the S-protein have the maximum contribution to the loss in binding affinity.

**Figure 2:**
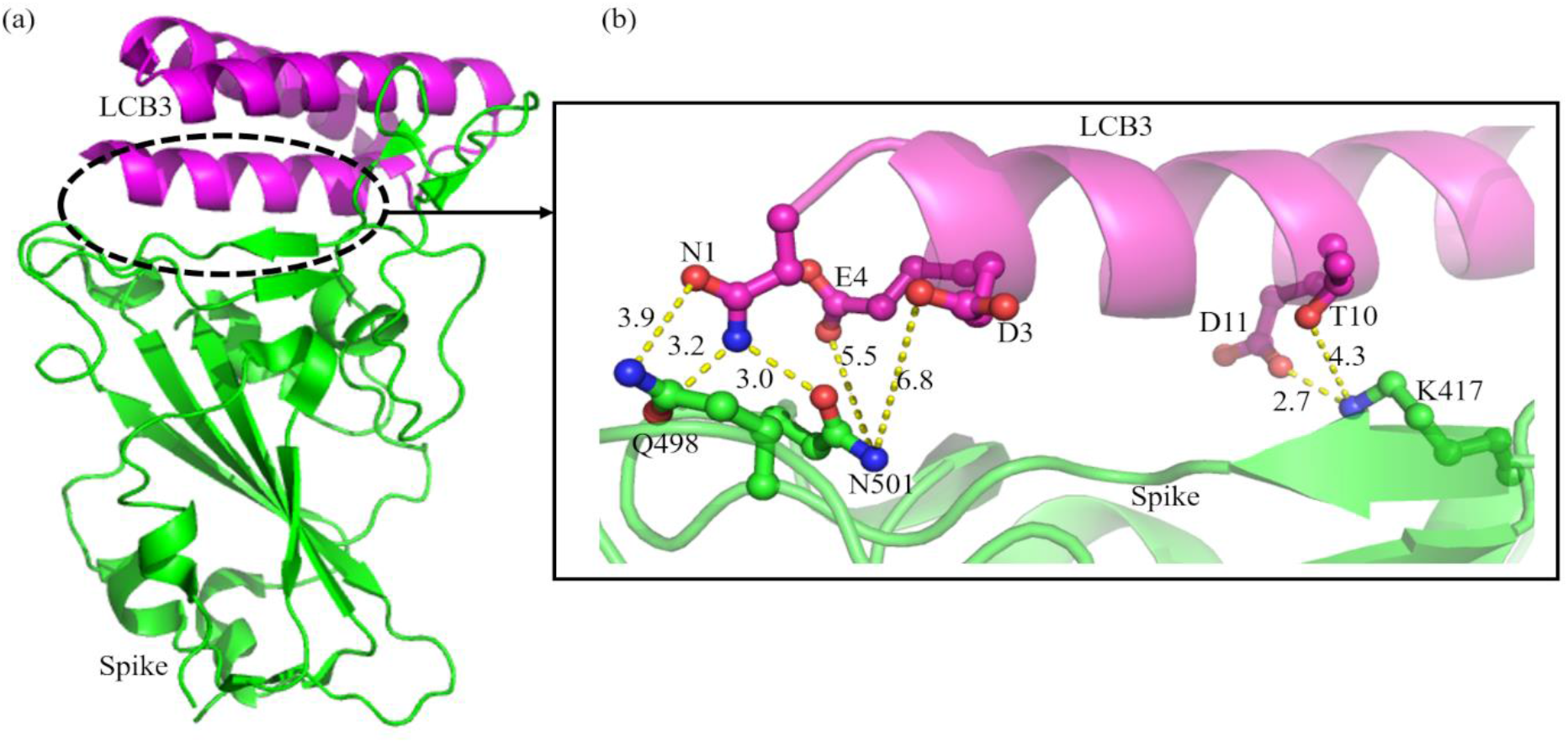
(a) Representative snapshot obtained after 10 ns simulation of PDB ID: 7JZM. (b) Detailed view of the binding interface: the interacting residues are shown in the stick representation. The distance among the oppositely charged atoms which may interact is given in Å.

**Table 2:**
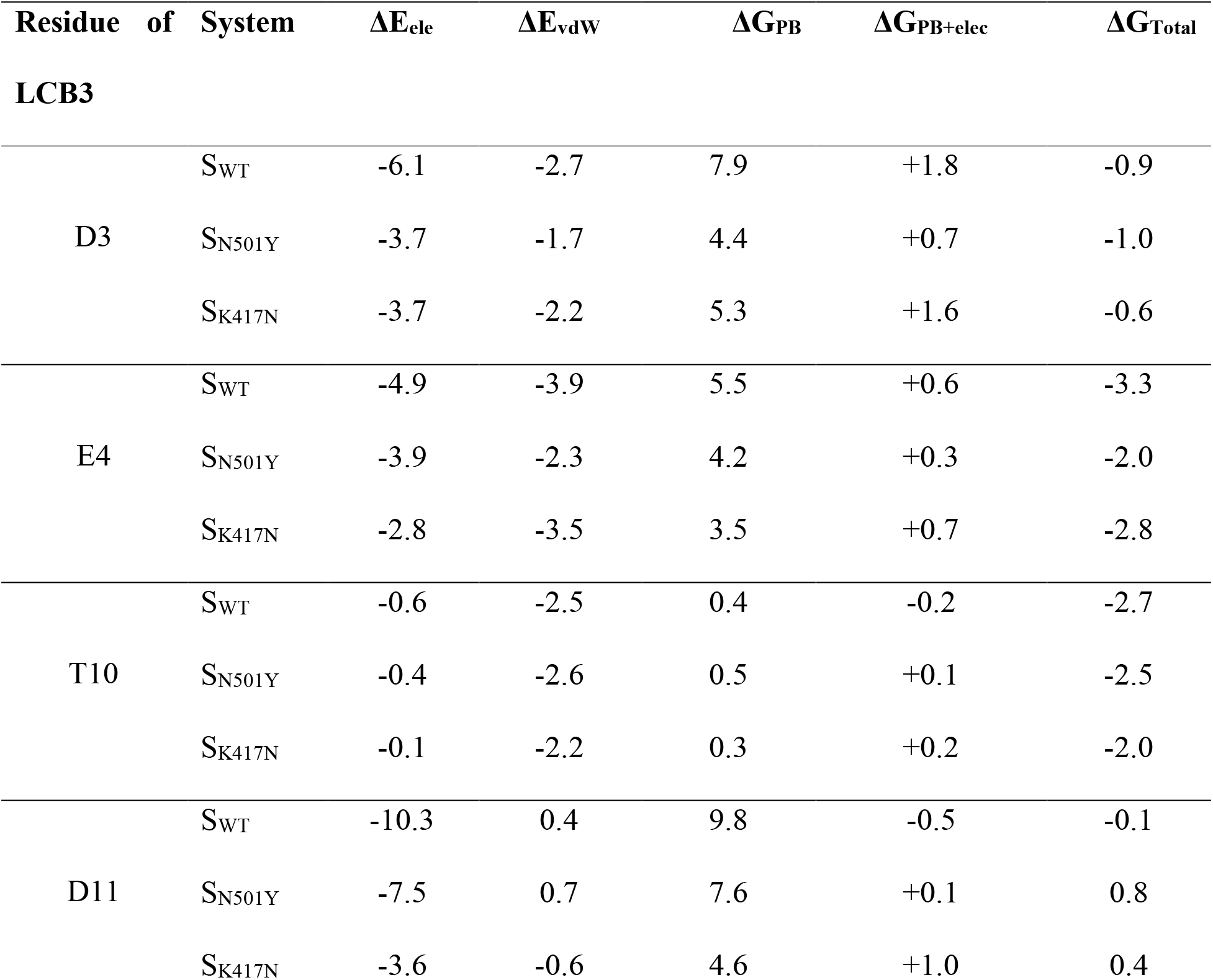
Free energy contributions of important residues of LCB3 present in the vicinity of N501 and K417, averaged over 1000 frames from the simulation of 10 independent trajectories each of 10 ns. Calculations were performed from the single trajectory method using the simulation of LCB3 complexed with wild-type and mutants of S-protein of SARS-CoV-2. All values are in kcal/mol.

### 3.3 Mutation Induced Conformational Changes in the Binding pattern of LCB3

Apart from the free energy contributions, it is also essential to understand the mutations induced conformational changes and interaction patterns to improve the binding affinity. In S_WT_ the N1, D3, and E4 of LCB3 form polar interaction with Q498 and N501 of S-protein (Figure 2 and 3a). The replacement of polar asparagine by hydrophobic tyrosine (N501Y) disrupts the electrostatic contribution (Figure 3b) of asparagine which was reflected in Table 1. In the structure of S-protein with ACE2 receptor, the polar residues are surrounded by two tyrosine residues Y41 of ACE2 and Y505 of S-protein (Figure 3c) ^92^ which was lacking with LCB3 inhibitor.

**Figure 3:**
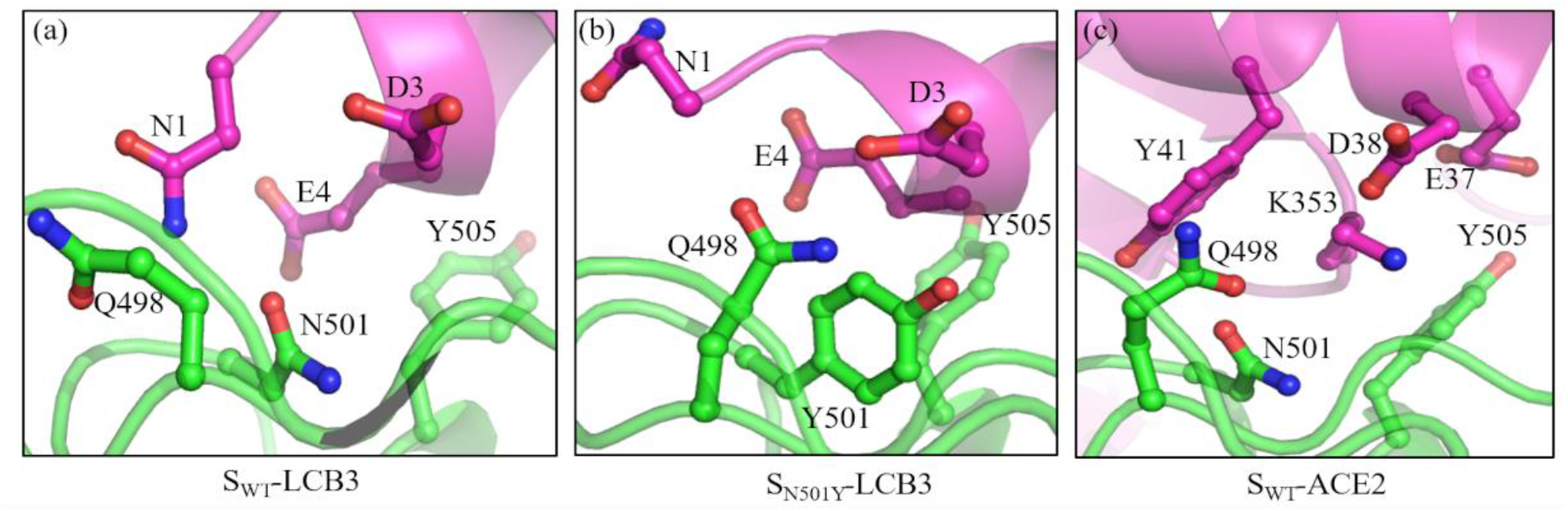
Interaction pattern in the binding interface of S-protein around N501 and the mutant Y501 (a) S_WT_ with LCB3 after 10 ns simulation of PDB ID:7JZM (b) S_N501Y_ with LCB3 after 10 ns simulation and (c) S_WT_ with ACE2; coordinates taken from PDB ID: 6M0J.

In S_K417N_, the replacement of a long side-chain residue lysine with asparagine weakens the interaction between S_K417N_ and LCB3 inhibitor (Figure 2, 4a, and 4b). This conformational change leads to the overall loss of +8.22 kcal/mol in the binding affinity of S_K417N_ with LCB3 inhibitor (Table 1).

**Figure 4:**
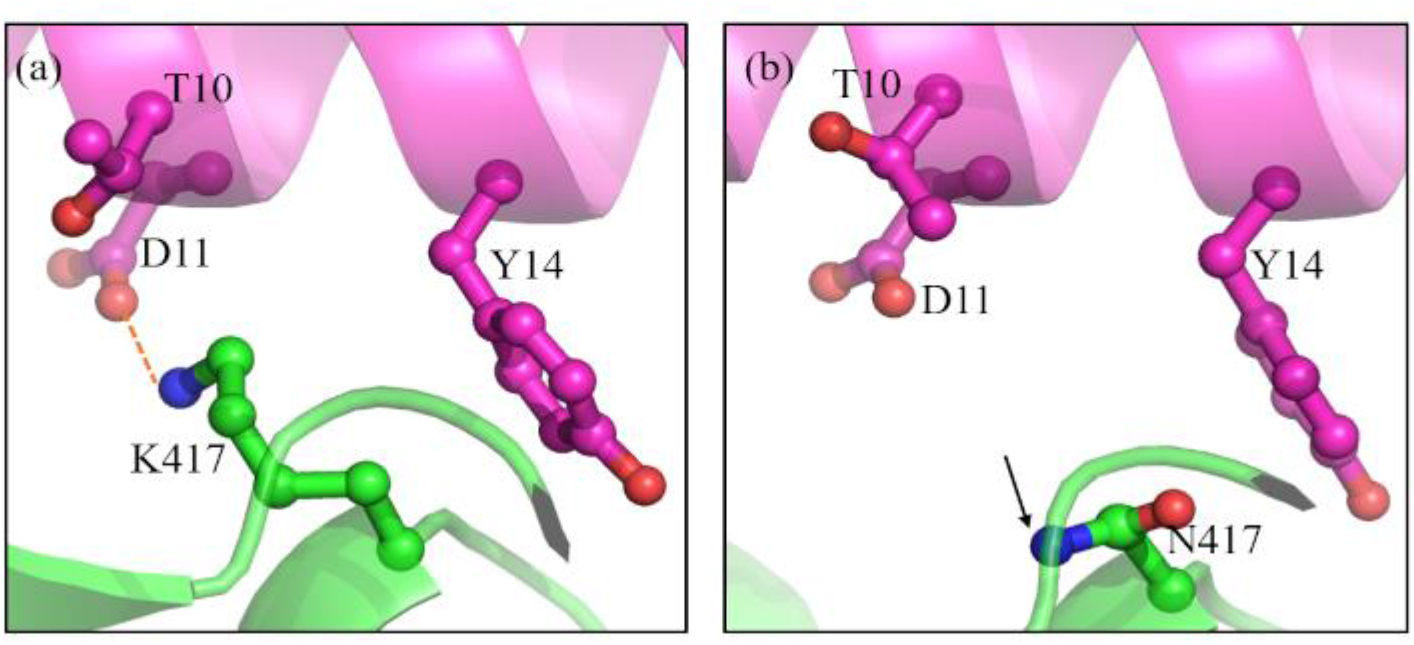
Representative snapshot obtained after 10 ns simulation of (a) S_WT_-LCB3: K417 of S-protein of SARS-CoV-2 is closer to D11 in S_WT_. (b) S_K417N_-LCB3: K417N mutation weakens the binding affinity of LCB3 with S_K417N_.

### 3.4. Substitution in LCB3 inhibitor for the improvement of the binding affinity with the mutants of S-protein

Based on the free energy and geometric analysis described in the previous section, we aimed to improve the interaction of Y501 and N417 of S_N501Y_ and S_K417N_ respectively with LCB3 by doing minimal change in LCB3. For this, the residues of LCB3 around Y501 of S_N501Y_ and K417 of S_K417N_ were targeted.

D3 and E4 of LCB3 were selected for modifications in S_N501Y_. There is a maximum of 19 possible mutations at each position, instead of testing them all here, we first tested the change of D3 by F and Y with the probability of forming pi-pi interactions or hydrophobic clusters with Y501 and Y505 (Figure 5a). Further, it may also help in improving the vdW contribution of Y3 of LCB3 which was lost by +1.0 kcal/mol in S_N501Y_ as compared to the S_WT_ (Table 2). MM-PBSA calculations show that the binding affinity of S_N501Y_-LCB3_D3F_ (−20.44 kcal/mol) and S_N501Y_-LCB3_D3Y_ (−23.68 kcal/mol) has improved as compared to the S_N501Y_-LCB3 (−15.69 kcal/mol). The improvement was due to the contribution of non-polar solvation energy (Table 1, 3). The effect of substitution of glutamic acid by tyrosine at the 4^th^ position of LCB3 was tested. The binding affinity of S_N501Y_-LCB3_E4Y_ was −18.25 kcal/mol, higher than the S_N501Y_-LCB3 but lower than the D3F and D3Y substitutions. The binding affinity of S_N501Y_-LCB3_D3Y_ (−23.68 kcal/mol) was similar to the S_WT_-LCB3 (−23.85 kcal/mol) therefore this modification can be successfully used against S_N501Y_. Further, we checked the binding affinity of LCB3_D3Y_ with S_WT_. Interestingly, we found that LCB3_D3Y_ has a higher affinity with S_WT_, −28.17 kcal/mol as compared to the LCB3 −23.85 kcal/mol (Table 1, 3). The increase in the affinity was due to the gain in vdW by −4.23 kcal/mol and polar contributions by −1.05 kcal/mol accompanied by loss of ~+1 kcal/mol in nonpolar components (Table 1, 3; Figure 5b).

**Figure 5:**
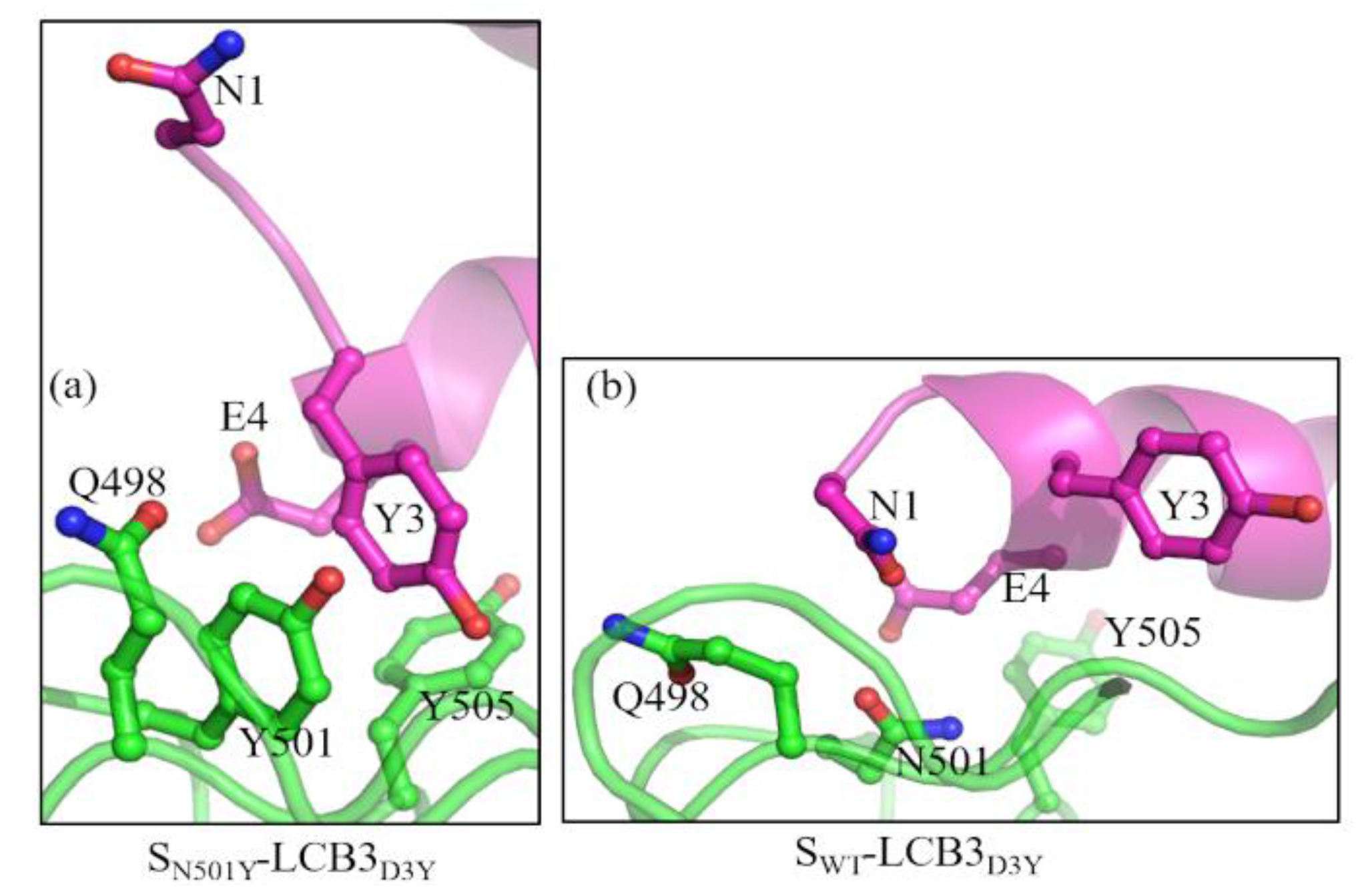
Representative conformation obtained after simulation of LCB3_D3Y_ with (a) S_N501Y_ (b) S_WT_. LCB3_D3Y_ allowed the formation of the hydrophobic cluster in S_N501Y_-LCB3_D3Y_.

**Table 3:**
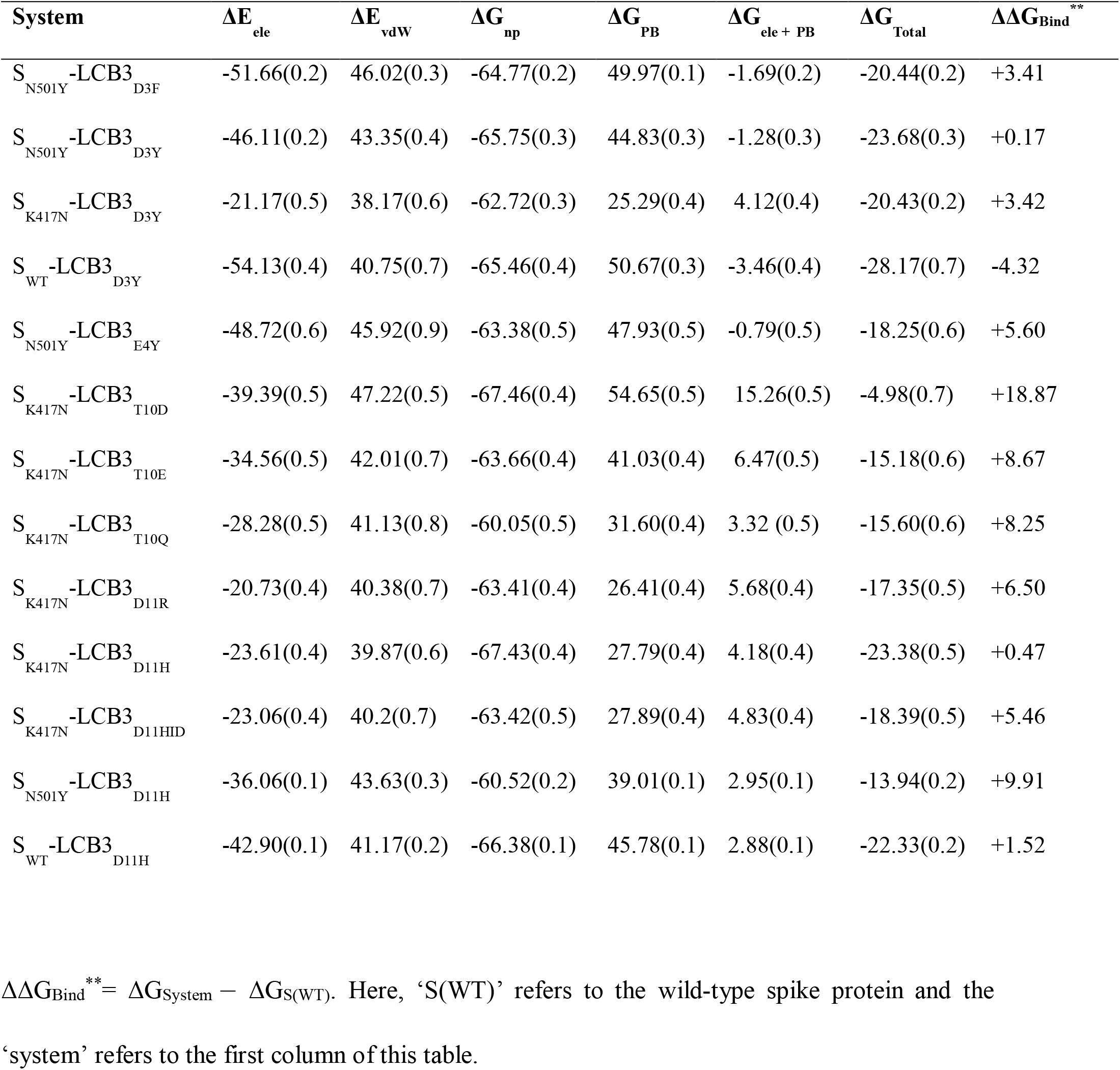
Binding free energy components of LCB3 variants with mutants and wild-type spike protein. Calculations averaged over 1000 frames obtained from 10 independent trajectories, each of 10 ns. The standard error is given in the parenthesis. Energy values are in kcal/mol.

| System | $\Delta E_{\text{ele}}$ | $\Delta E_{\text{vdW}}$ | $\Delta G_{\text{np}}$ | $\Delta G_{\text{PB}}$ | $\Delta G_{\text{ele + PB}}$ | $\Delta G_{\text{Total}}$ | $\Delta\Delta G_{\text{Bind}}^{**}$ |
| --- | --- | --- | --- | --- | --- | --- | --- |
| S<sub>N501Y</sub>-LCB3<sub>D3F</sub> | -51.66(0.2) | 46.02(0.3) | -64.77(0.2) | 49.97(0.1) | -1.69(0.2) | -20.44(0.2) | +3.41 |
| S<sub>N501Y</sub>-LCB3<sub>D3Y</sub> | -46.11(0.2) | 43.35(0.4) | -65.75(0.3) | 44.83(0.3) | -1.28(0.3) | -23.68(0.3) | +0.17 |
| S<sub>K417N</sub>-LCB3<sub>D3Y</sub> | -21.17(0.5) | 38.17(0.6) | -62.72(0.3) | 25.29(0.4) | 4.12(0.4) | -20.43(0.2) | +3.42 |
| S<sub>WT</sub>-LCB3<sub>D3Y</sub> | -54.13(0.4) | 40.75(0.7) | -65.46(0.4) | 50.67(0.3) | -3.46(0.4) | -28.17(0.7) | -4.32 |
| S<sub>N501Y</sub>-LCB3<sub>E4Y</sub> | -48.72(0.6) | 45.92(0.9) | -63.38(0.5) | 47.93(0.5) | -0.79(0.5) | -18.25(0.6) | +5.60 |
| S<sub>K417N</sub>-LCB3<sub>T10D</sub> | -39.39(0.5) | 47.22(0.5) | -67.46(0.4) | 54.65(0.5) | 15.26(0.5) | -4.98(0.7) | +18.87 |
| S<sub>K417N</sub>-LCB3<sub>T10E</sub> | -34.56(0.5) | 42.01(0.7) | -63.66(0.4) | 41.03(0.4) | 6.47(0.5) | -15.18(0.6) | +8.67 |
| S<sub>K417N</sub>-LCB3<sub>T10Q</sub> | -28.28(0.5) | 41.13(0.8) | -60.05(0.5) | 31.60(0.4) | 3.32 (0.5) | -15.60(0.6) | +8.25 |
| S<sub>K417N</sub>-LCB3<sub>D11R</sub> | -20.73(0.4) | 40.38(0.7) | -63.41(0.4) | 26.41(0.4) | 5.68(0.4) | -17.35(0.5) | +6.50 |
| S<sub>K417N</sub>-LCB3<sub>D11H</sub> | -23.61(0.4) | 39.87(0.6) | -67.43(0.4) | 27.79(0.4) | 4.18(0.4) | -23.38(0.5) | +0.47 |
| S<sub>K417N</sub>-LCB3<sub>D11HID</sub> | -23.06(0.4) | 40.2(0.7) | -63.42(0.5) | 27.89(0.4) | 4.83(0.4) | -18.39(0.5) | +5.46 |
| S<sub>N501Y</sub>-LCB3<sub>D11H</sub> | -36.06(0.1) | 43.63(0.3) | -60.52(0.2) | 39.01(0.1) | 2.95(0.1) | -13.94(0.2) | +9.91 |
| S<sub>WT</sub>-LCB3<sub>D11H</sub> | -42.90(0.1) | 41.17(0.2) | -66.38(0.1) | 45.78(0.1) | 2.88(0.1) | -22.33(0.2) | +1.52 |

$\Delta\Delta G_{\text{Bind}}^{**} = \Delta G_{\text{System}} - \Delta G_{\text{S(WT)}}$ . Here, ‘S(WT)’ refers to the wild-type spike protein and the ‘system’ refers to the first column of this table.

To overcome the loss in the binding affinity of S_K417N_, two residues of LCB3: T10, and D11 were targeted for substitutions, based on the loss in the free energy contributions of the residues, and the distance from the mutated K417 (Table 2, Figure 2 and 4). Substitutions of the amino acid T10 of LCB3 by D, E and Q were tested with the possibility to improve polar interaction with N417 but none of these have enhanced the affinity (Table 3). D11 is closer to K417 than T10 (Figure 2), forms direct polar interaction with K417 (Figure 2b), and there is a loss of +0.5 kcal/mol in the free energy contributions of D11 in S_K417N_ as compared to S_WT_ (Table 2). Therefore, D11 was replaced with a bulky and long side-chain amino acid that can interact with N417. In this regard, two substitutions, D11R and D11H were tested. The binding affinity of the LCB3_D11R_ with K417N is −17.35 kcal/mol, ~+6.5 kcal/mol lesser affinity than the S_WT_-LCB3 (Table 3). However, LCB3_D11H_ showed a higher binding affinity (−23.38 kcal/mol) with S_K417N_ (Table 3). Here, two different protonation states of histidine, at ND1 (HID) and HE2 (HIE) were tested. Histidine with protonation at HE2 (HIE) has shown higher binding affinity as compared to HID. ‘H’ refers to the ‘HIE’ protonation state in this paper. From the structure, it was found that the imidazole ring moves towards the binding interface and comes closer to N417 (Figure 6 a), and strengthens the binding by improving the nonpolar solvation component (Table 2). The binding affinity of LCB3_D11H_ with S_WT_ is −22.33 kcal/mol, slightly less than the binding affinity of LCB3. In S_WT_-LCB3_D11H_, K417 of S-protein repels the H11 slightly away from the binding interface (Figure 6b).

**Figure 6:**
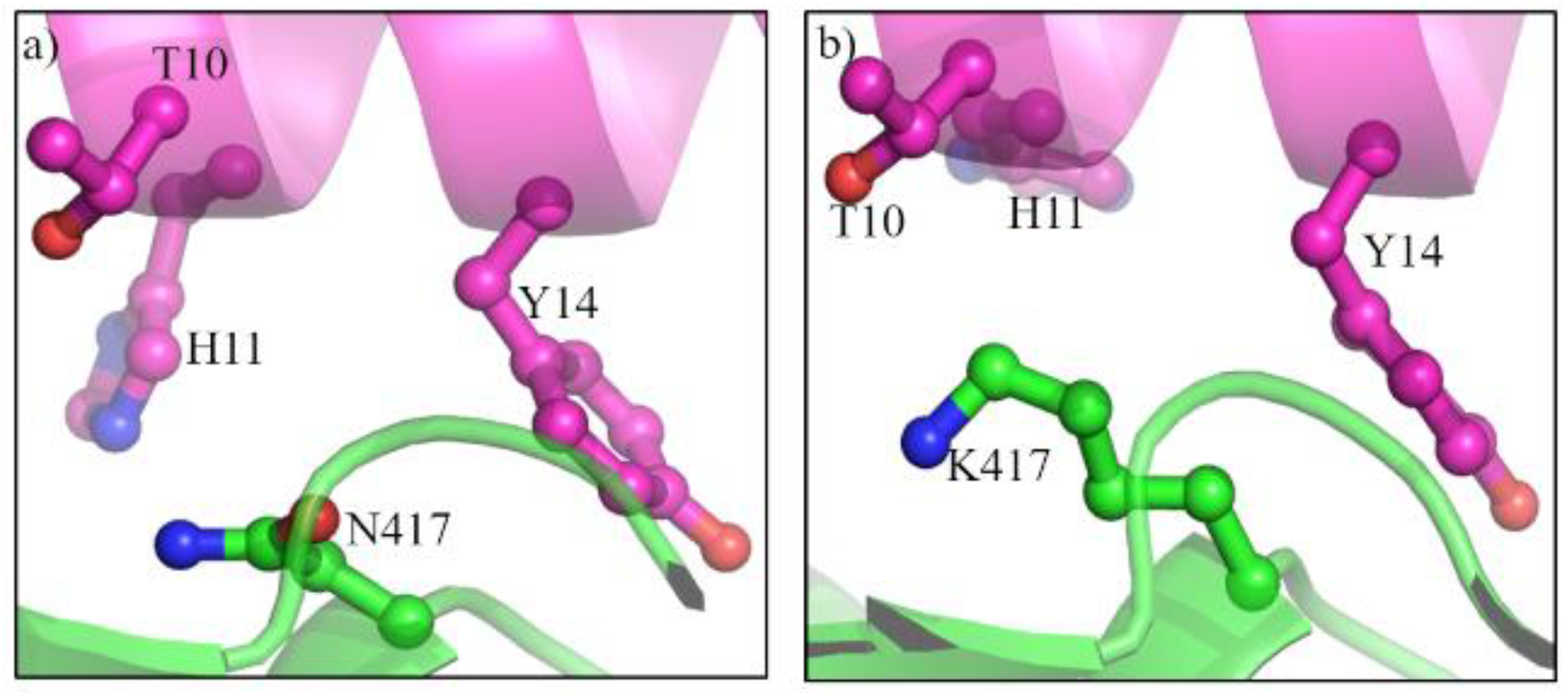
Representative snapshot of (a) S_K417N_-LCB3_D11H_; substituted histidine (H11) makes the interaction more compact with S_K417N_ as compared to the LCB3 in Figure 4b. (b) S_WT_-LCB3_D11H_; the long side chain of K417 pushed H11 away from the binding interface.

Further, the binding affinity of S_K417N_ with LCB3_D3Y_ was tested, which has already shown good affinity with S_WT_ and S_N501Y_. The binding affinity of S_K417N_-LCB3_D3Y_ was found −20.43 kcal/mol.

It showed that the LCB3_D3Y_ can bind efficiently with the S_WT_ (−28.17), S_N501Y_ (−23.68 kcal/mol), and also with S_K417N_ (−20.43 kcal/mol) when compared to the original LCB3 inhibitor. Further, S_K417N_ has a stronger binding affinity with LCB3_D11H_ (−23.38 kcal/mol) as compared to the LCB3_D3Y_ (−20.43kcal/mol). However, LCB3_D11H_ does not bind efficiently with S_N501Y_ (−13.95 kcal/mol).

In the binding free energy calculations described so far, the entropy of solute is not considered, a common practice in MM-PBSA calculations. However, for the present problem, the calculation of solute entropy is important as changes in amino acids in either S-protein or LCB3 can change the flexibility of the system leading to a change in entropy. As the entropy calculations are computationally intensive, we have performed it for some important systems as mentioned in Table 4. The change in the entropy was calculated over separate 1μs trajectories complex, receptor and ligand. The entropy essentially converges for each system in 1μs MD simulation (Figure 7a to c). The negative of the change in entropy multiplied by temperature, T (the last term of equation (3), -TΔS) of LCB3 (+55.5 kcal/mol) and its two variants LCB3_D3Y_ (+56.0 kcal/mol) and LCB3_D11H_ (+56.2 kcal/mol) with S_WT_ were almost similar with no significant differences between them (Figure 7d, Table S4). However, the significant differences in -TΔS were observed for S_N501Y_-LCB3_D3Y_ (+49.4 kcal/mol) and S_K417N_-LCB3_D11H_ (+24.4 kcal/mol). In both these cases, the entropy change on binding was significantly reduced. This reduction in the change of entropy will also help in improving the binding affinity of these variants of LCB3 with mutated S-protein (equation 3). It reflects that the binding affinity of S_WT_ with the proposed variants of LCB3 is mainly enthalpy driven; however, in the mutated S-protein (S_N501Y_ and S_K417N_) entropic changes played significant contributions in improving the binding affinity (Table 4).

**Table 4:** Entropy Change over 1 μs simulation of the LCB3 and its successful variants with the wild-type and mutated S-protein. The values of negative change in entropy multiplied by T (-TΔS) is in kcal/mol.

| System | Entropy (-TΔS) over 1 μs simulation |
| --- | --- |
| S<sub>WT</sub>-LCB3 | +55.5 |
| S<sub>N501Y</sub>-LCB3<sub>D3Y</sub> | +49.4 |
| S<sub>K417N</sub>-LCB3<sub>D11H</sub> | +24.4 |
| S<sub>WT</sub>-LCB3<sub>D3Y</sub> | +56.0 |
| S<sub>WT</sub>-LCB3<sub>D11H</sub> | +56.2 |

**Figure 7:**
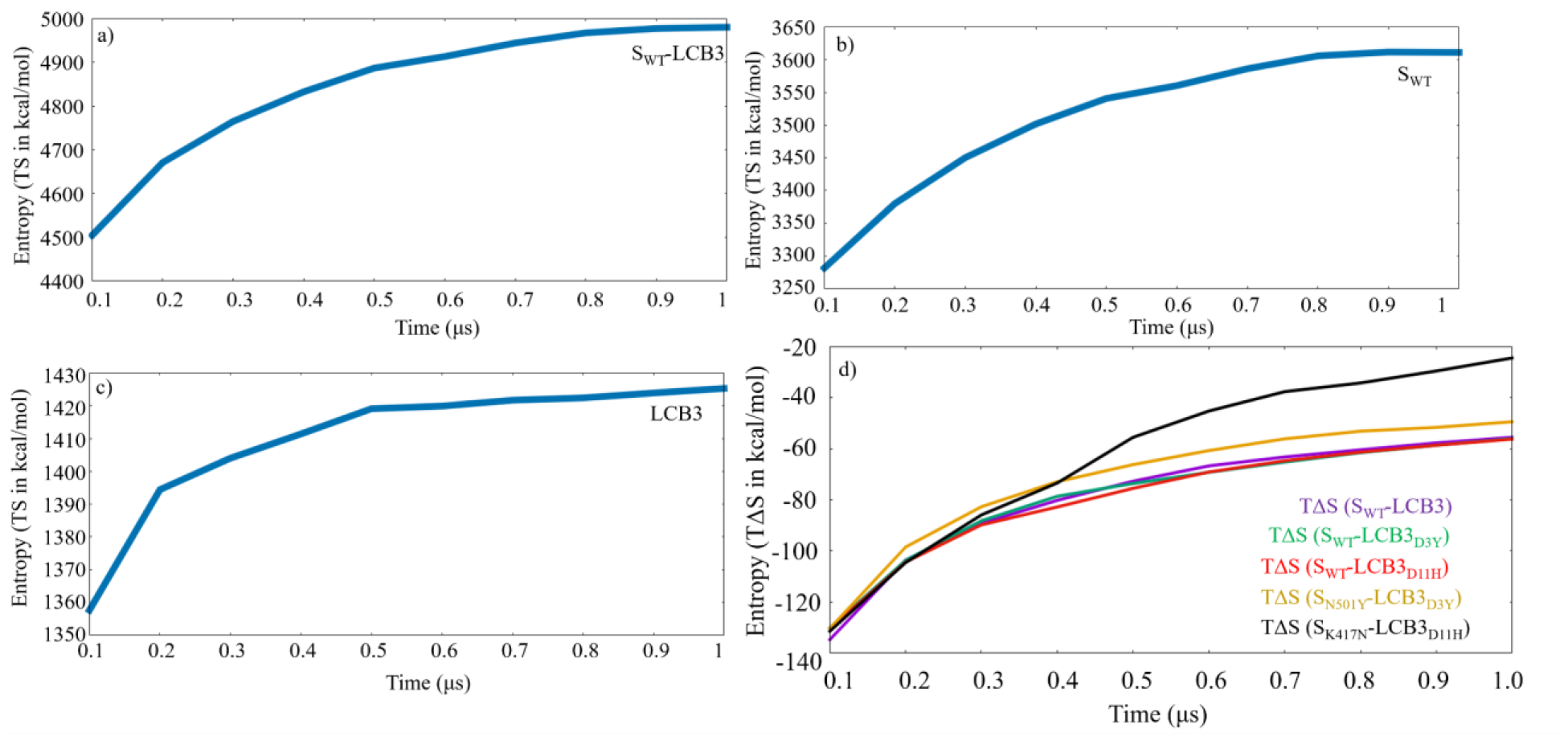
Convergence plot of (a) TS over 1 μs simulation of S_WT_-LCB3 (b) S_WT_ (c) LCB3 (d) TΔS of LCB3 and its proposed variant LCB3_D3Y_ and LCB3_D11H_ with S_WT_, S_N501Y_ and S_K417N_ over 1 μs simulation.

To summarize, our calculation and analysis show that a judicious combination of free energy decomposition and geometric consideration can suggest ways to improve the inhibitor against specific mutations. Detailed binding affinity calculation that includes solute entropy shows that even a single amino acid change in LCB3 can make it a potent inhibitor to the S-protein and its mutants. In particular, LCB3_D3Y_ and LCB3_D11H_ were proposed as the variants of LCB3 that may inhibit the WT as well as mutants of S-protein. It is to be noted that LCB3 is taken as a prototype for more complex systems such as antibodies. Moreover, this methodology is general so that it can be applied to any inhibitor against different targets.

Our calculation methodology does have some caveats. First, as we wanted to develop a fast methodology, only the receptor-binding domain of the S-protein and the LCB inhibitor were considered. The ACE-2 protein is not considered in the calculation. Our choice of using a simpler system is similar to the work done by williams et al ^90^.

## 4. Conclusions

The fight against SARS-CoV-2 requires further development of vaccines and medicines. One of the main targets to develop a drug against this virus is the spike protein of the virus. Although there are promising drug candidates against the virus, the major issue is how to design/modify inhibitors against the various mutant strains of the virus. In this work, as a proof-of-principle study, we have considered a mini-protein inhibitor, LCB3 that binds to the spike protein of SARS-CoV-2. We have devised a computational protocol to modify LCB3 against the two most common mutations in the spike protein, namely, N501Y and K417N. Our computational methodology includes a free energy decomposition procedure in conjunction with a detailed analysis of the binding site. Our proposed modified LCB3 is shown to bind to the two mutant forms of the spike protein potently. The binding enthalpy showed that the single residue mutations LCB3_D3Y_ work well against S_WT_ as well as S_N501Y_ and S_K417N_. Another modification LCB3_D11H_ binds efficiently with S_K417N_ and S_WT_ but not with S_N501Y_. Further, the entropic contributions also favor the binding of LCB3_D3Y_ with S_N501Y_, and LCB3_D11H_ with S_K417N_ respectively. The study found that LCB3_D3Y_ and LCB3_D11H_ were bound with a higher affinity with the WT as well as mutated S-protein. This strategy can be useful to redesign the peptide-based inhibitor against the target protein that undergoes frequent mutation.

## Supporting information

https://drive.google.com/file/d/1TL6-N2wgC_2p3G4pH1nXGUof3YFGgOb0/view?usp=sharing

