## Supplementary material for "A strategy to optimize the peptide-based inhibitors against different mutants of the spike protein of SARS-CoV-2": https://drive.google.com/file/d/1TL6-N2wgC_2p3G4pH1nXGUof3YFGgOb0/view?usp=sharing


### Supporting Information

1. **Table S1:** List of the simulation performed for the MM-PBSA calculation using single trajectory method.
2. **Table S2:** List of the simulation performed for estimation of the change in the Entropy.
3. **Table S3:** Test to select the dielectric constant of solute.
4. **Table S4:** Change in the entropy obtained from separate simulation of complex, receptor, and ligand over 1 μs simulation.

**Table S1:** List of the simulated system with total sampling time used for the MM-PBSA calculations by single trajectory method

| System | Total Simulation time (Simulation Length × No. of Independent Simulation) |
|---|---|
| $S_{N501Y}$-LCB3$_{D3F}$ | 100 ns (10 ns ×10 ) |
| $S_{N501Y}$-LCB3$_{D3Y}$ | 100 ns (10 ns ×10 ) |
| $S_{K417N}$-LCB3$_{D3Y}$ | 100 ns (10 ns ×10 ) |
| $S_{WT}$-LCB3$_{D3Y}$ | 100 ns (10 ns ×10 ) |
| $S_{N501Y}$-LCB3$_{E4Y}$ | 100 ns (10 ns ×10 ) |
| $S_{K417N}$-LCB3$_{T10D}$ | 100 ns (10 ns ×10 ) |
| $S_{K417N}$-LCB3$_{T10E}$ | 100 ns (10 ns ×10 ) |
| $S_{K417N}$-LCB3$_{T10Q}$ | 100 ns (10 ns ×10 ) |
| $S_{K417N}$-LCB3$_{D11R}$ | 100 ns (10 ns ×10 ) |
| $S_{K417N}$-LCB3$_{D11H}$ | 100 ns (10 ns ×10 ) |
| $S_{K417N}$-LCB3$_{D11HID}$ | 100 ns (10 ns ×10 ) |
| $S_{N501Y}$-LCB3$_{D11H}$ | 100 ns (10 ns ×10 ) |
| $S_{WT}$-LCB3$_{D11H}$ | 100 ns (10 ns ×10 ) |

**Table S2:** List of the separate simulation of complex, receptor and ligand for quasi-harmonic approximation.

| System | Simulation Length |
|---|---|
| $S_{WT}$ | 1μs |
| $S_{N501Y}$ | 1μs |
| $S_{K417N}$ | 1μs |
| LCB3 | 1μs |
| $S_{WT}$-LCB3 | 1μs |
| $LCB3_{D3Y}$ | 1μs |
| $LCB3_{D11H}$ | 1μs |
| $S_{N501Y}$- $LCB3_{D3Y}$ | 1μs |
| $S_{K417N}$- $LCB3_{D11H}$ | 1μs |
| $S_{WT}$- $LCB3_{D3Y}$ | 1μs |
| $S_{WT}$- $LCB3_{D11H}$ | 1μs |
| Total | 11 μs |

**Table S3:** Change in the binding free energy tested at the dielectric constant of protein 1, 2, 4 and 8. Values of binding energy are in kcal/mol. The calculations were performed over 10 trajectories each of 10 ns. 1000 frames were used in the calculations. Standard error is given in small parenthesis.

| System | Dielectric Constant | Binding Energy |
|---|---|---|
| 1. $S_{WT}$-LCB3[1] | 1 | +8.28 (1.4) |
| | 2 | -9.61 (0.8) |
| | 4 | -18.90 (0.6) |
| | 8 | -23.85 (0.6) |
| 2. $S_{N501Y}$-LCB3[2] | 1 | +25.63 (2.0) |
| | 2 | +2.34 (1.0) |
| | 4 | -9.53 (0.6) |
| | 8 | -15.69 (0.6) |
| | 1 | +17.35 |
| 3. Difference (2-1) | 2 | +11.95 |
| | 4 | +9.37 |
| | 8 | +8.16 |

Change in the binding affinity was tested using dielectric constants 1, 2, 4 and 8 for the protein. It was observed that the trend of the binding affinity of $S_{WT}$-LCB3 and $S_{N501Y}$-LCB3 remains the same in all the calculations i.e, the binding affinity of $S_{N501Y}$ was reduced with LCB3 as compared to $S_{WT}$. The difference in the binding affinity at dielectric constant 8 is more similar (~+8.1 kcal/mol) to the already reported value (+7.1 kcal/mol) by Williams et al.[1]. Therefore, the dielectric constant of protein, 8 was selected for all the calculations.

**Table S4:** Change in Entropy at a given time, obtained from the separate, 1μs simulation of complex, receptor and ligand. Calculations were performed using 5 lakhs frames. TΔS was obtained by the subtraction of TS of receptor and ligand from the complex.

| System | Time (μs) | TΔS (kcal/mol) |
|---|---|---|
| | 0.1 | -134.8 |
| | 0.2 | -103.6 |
| | 0.3 | -89.2 |
| | 0.4 | -80.2 |
| $S_{WT}$-LCB3 | 0.5 | -72.7 |
| | 0.6 | -66.7 |
| | 0.7 | -63.2 |
| | 0.8 | -60.4 |
| | 0.9 | -57.7 |
| | 1.0 | **-55.5** |
| | 0.1 | -130.4 |
| | 0.2 | -98.5 |
| | 0.3 | -82.7 |
| | 0.4 | -72.9 |
| | 0.5 | -66.2 |
| $S_{N501Y}$-$LCB3_{D3Y}$ | 0.6 | -60.7 |
| | 0.7 | -56.1 |
| | 0.8 | -53.1 |
| | 0.9 | -51.6 |
| | 1.0 | **-49.4** |
| | 0.1 | -130.2 |
| | 0.2 | -103.7 |
| | 0.3 | -88.2 |
| | 0.4 | -78.6 |

| | | |
|---|---|---|
| $S_{WT}$-LCB3$_{D3Y}$ | 0.5 | -73.6 |
| | 0.6 | -69.2 |
| | 0.7 | -65.2 |
| | 0.8 | -61.5 |
| | 0.9 | -58.6 |
| | 1.0 | **-56.0** |
| $S_{K417N}$-LCB3$_{D11H}$ | 0.1 | -131.5 |
| | 0.2 | -104.6 |
| | 0.3 | -85.9 |
| | 0.4 | -73.4 |
| | 0.5 | -55.5 |
| | 0.6 | -45.2 |
| | 0.7 | -37.6 |
| | 0.8 | -34.2 |
| | 0.9 | -29.5 |
| | 1.0 | **-24.4** |
| | 0.1 | -130.5 |
| | 0.2 | -104.4 |
| | 0.3 | -89.8 |
| | 0.4 | -82.8 |
| $S_{WT}$-LCB3$_{D11H}$ | 0.5 | -75.5 |
| | 0.6 | -69.1 |
| | 0.7 | -64.7 |
| | 0.8 | -61.3 |
| | 0.9 | -58.6 |
| | 1.0 | **-56.2** |
